## supplementary figures for "*Tgfbr1* regulates lateral plate mesoderm and endoderm reorganization during the trunk to tail transition"

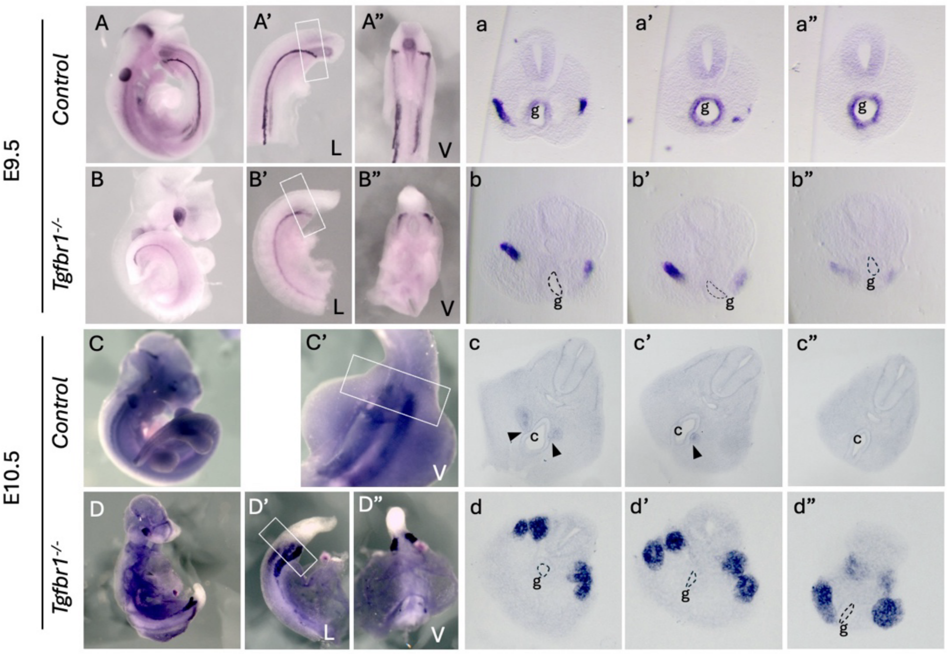


**Supplementary figure 1. Expression of the IM marker *Pax2* in *Tgdfr1*$^{-/-}$ embryos. (**A-b") *Pax2* expression in E9.5 control (A-a") and *Tgfbr1*$^{-/-}$ (B-b") embryos. (a-a") and (b-b") show sections through the region marked by the rectangle in A' and B'. **(**C-d") *Pax2* expression in E10.5 control (C-c") and *Tgfbr1*$^{-/-}$ (D-d") embryos. (c-c") and (d-d") show sections through the region marked by the rectangle in C' and D'. At E10.5 pronephric ducts merge with the cloaca in control embryos (black arrowheads in c, c'), but not in the mutants (d, d"). c- cloaca, g- gut, L – lateral, V - ventral.

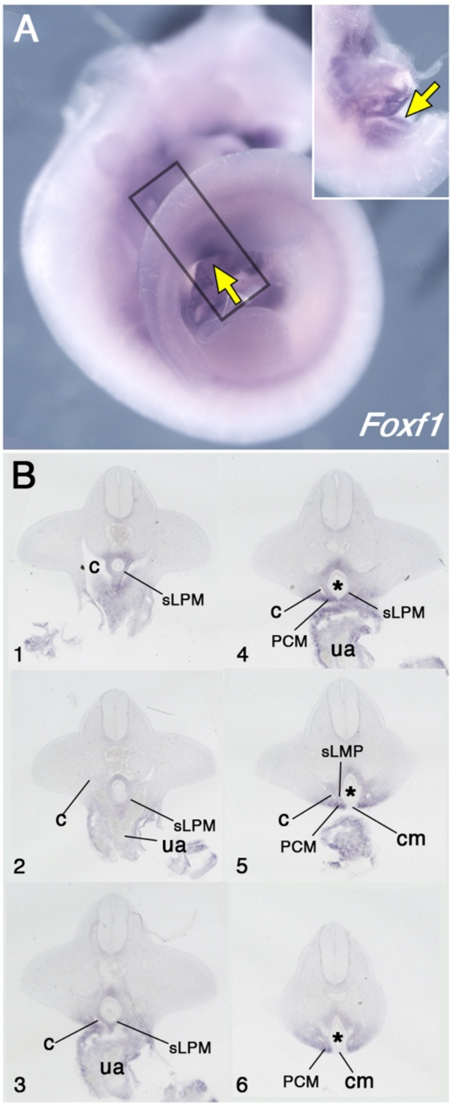


**Supplementary figure 2. Pericloacal mesenchyme derives from the mesoderm adjacent to the allantois. A.** Whole mount in situ hybridization hybridizations showing expression of *Foxf1* in E10.5 wild type embryo. Yellow arrows show expression in the pericloacal mesenchyme. Inset shows ventral view in the pericloacal mesenchyme. **B.** Series of transversal sections through the region marked by the rectangle in A (1-6, from anterior to posterior). Splanchnic LPM (sLPM) is separated from the pericloacal mesenchyme (PCM) by the coelomic cavity (c). Cloaca is labelled by asterisk, ua – umbilical artery, cm – cloaca membrane.

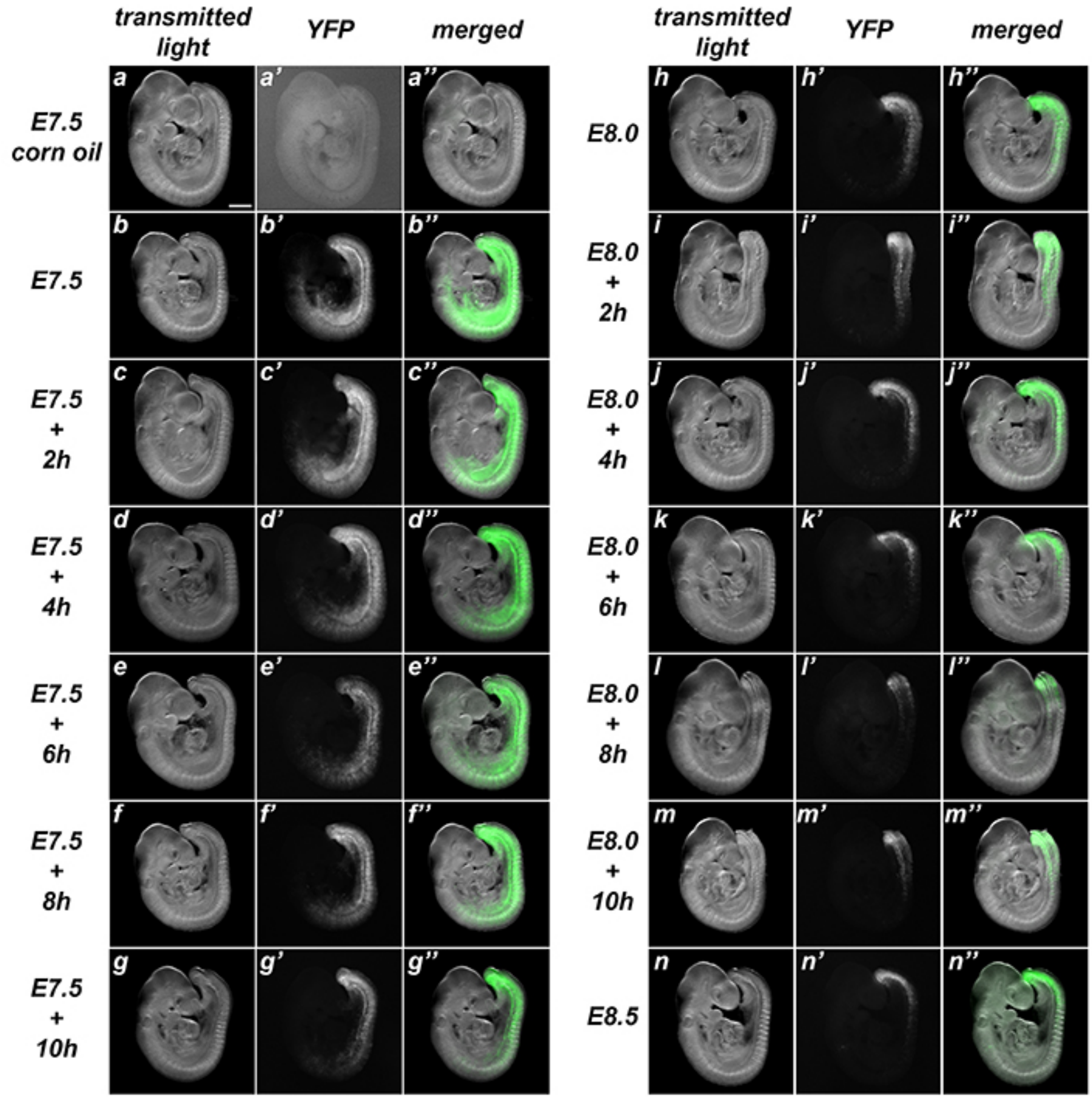


**Supplementary figure 3. Characterization of the recombination activity of the *Tstr-cre*$^{ERT}$ transgenics.** These transgenics were analyzed by crossing them with *ROSA26R-YFP* mice. While non-treated embryos showed only rare events of spontaneous recombination (a), administration of tamoxifen at early stages induced extensive recombinant territories that got progressively restricted to the caudal region as tamoxifen was being administered at later time-points (b-n). Left rows adjacent to images show the time of tamoxifen administration. Size bar: 200 μm

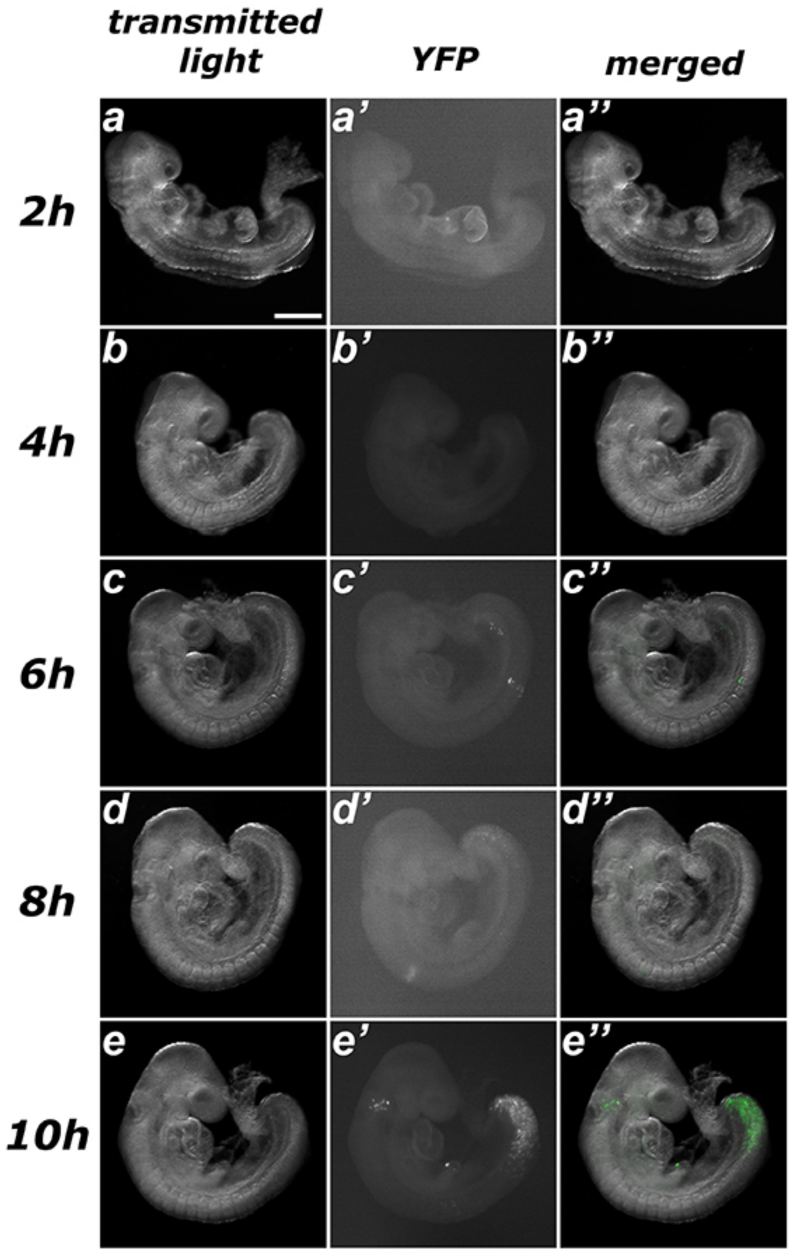


**Supplementary figure 4. Estimating the time required for recombination after tamoxifen administration in *Tstr-cre$^{ERT}$::ROSA26R-YFP* embryos.** No evident sign of recombination was observed after up to six hours of treatment (a-c), only a few scarce spontaneous events. Embryos harvested eight hours after tamoxifen administration exhibited early signs of recombination (d) and clear induction was observed ten hours after treatment (e). Size bar: 200 μm.

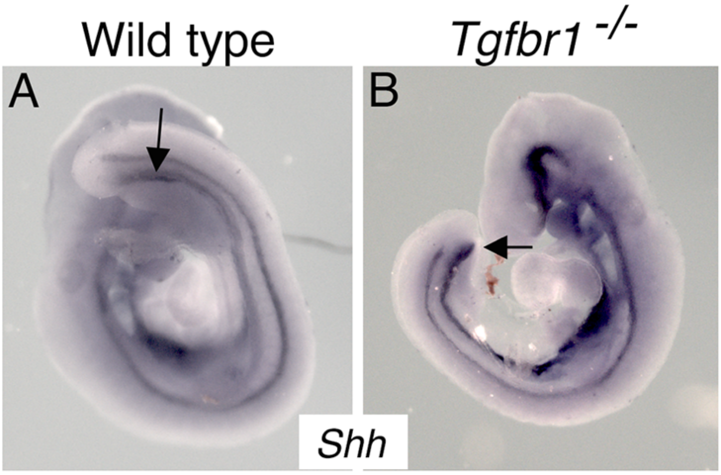


**Supplementary Figure 5.** Whole mount in situ hybridization on E9.5 wild type (A) and *Tgfbr1*$^{-/-}$ (B) embryos with a probe for *Shh*. In wild type embryos, the endoderm will form the cloaca at the level of the developing hindlimb (arrow) and extend into the emerging tail bud. In the mutant embryo the endoderm finishes at the posterior embryonic end merging into the ventral wall of the embryo (arrow).

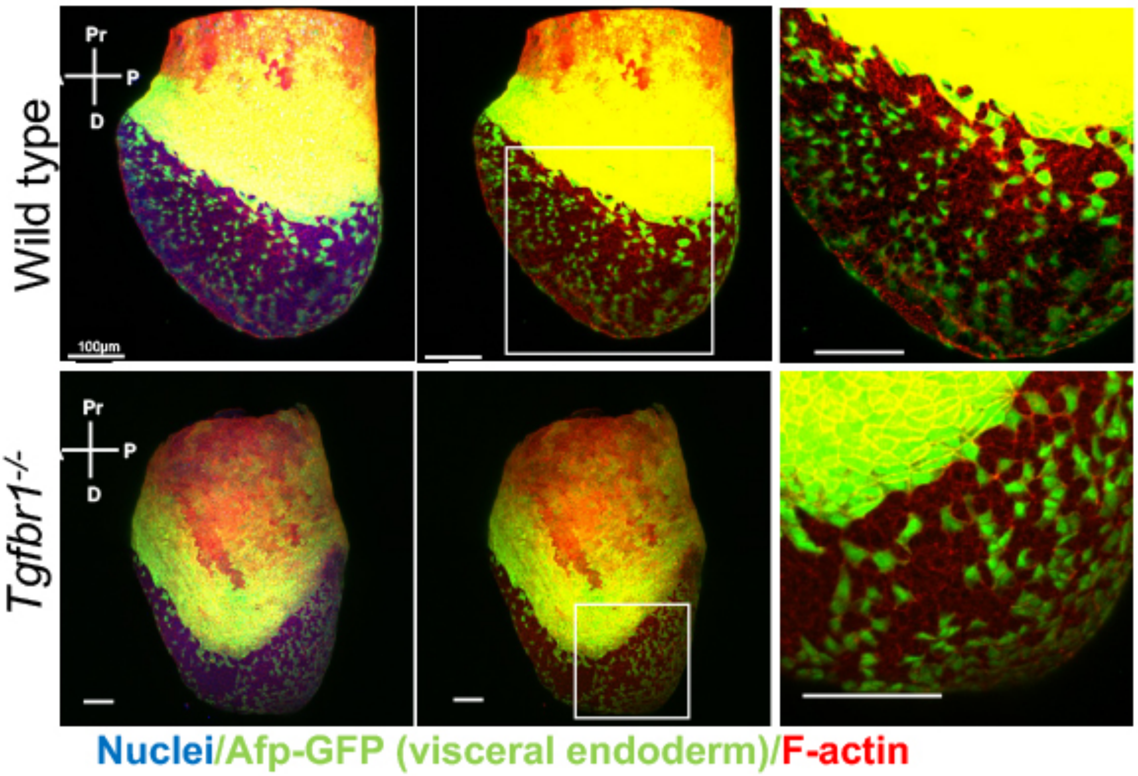


**Supplementary Figure 6.** Analysis of visceral endoderm (VE) dispersal in wild type and *Tgfbr1* mutant E7.5 embryos. No differences can be seen in the mutant embryo relative to the wild type control.

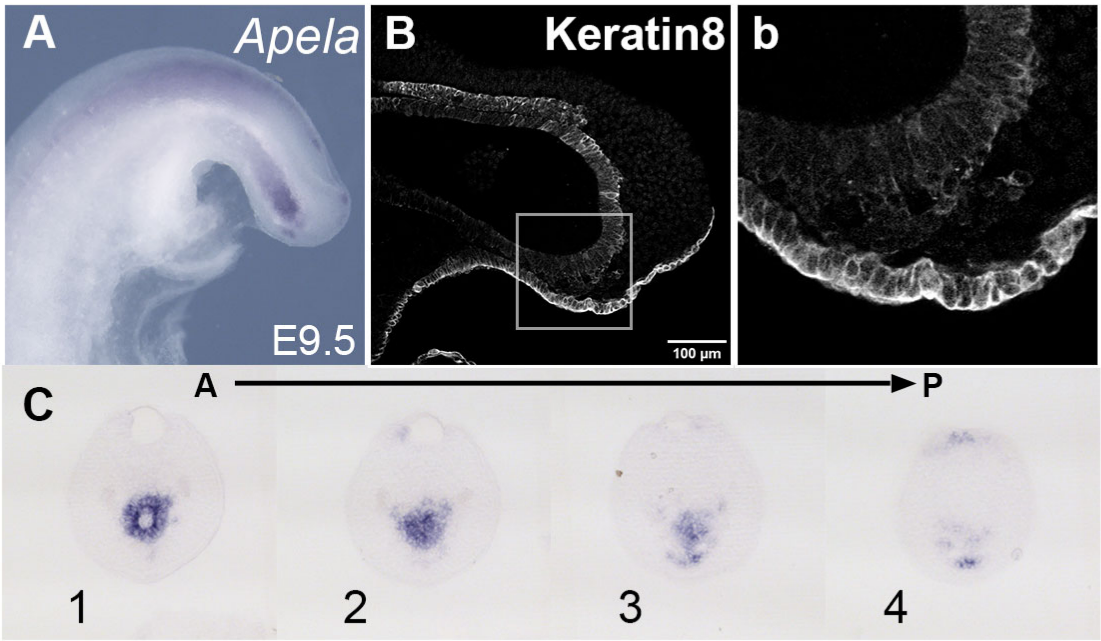


**Supplementary figure 7. Tail gut of E9,5 *wild type* embryo. A, C**. *Apela* staining in the wild type tail bud at E9.5. C shows a series of transversal sections through the Apela-positive region shown in the whole mount image in A. **B, b.** Keratin 8 stained cells ventrally and posteriorly to the tail gut endoderm. b shows magnified image of the region marked by square in B.

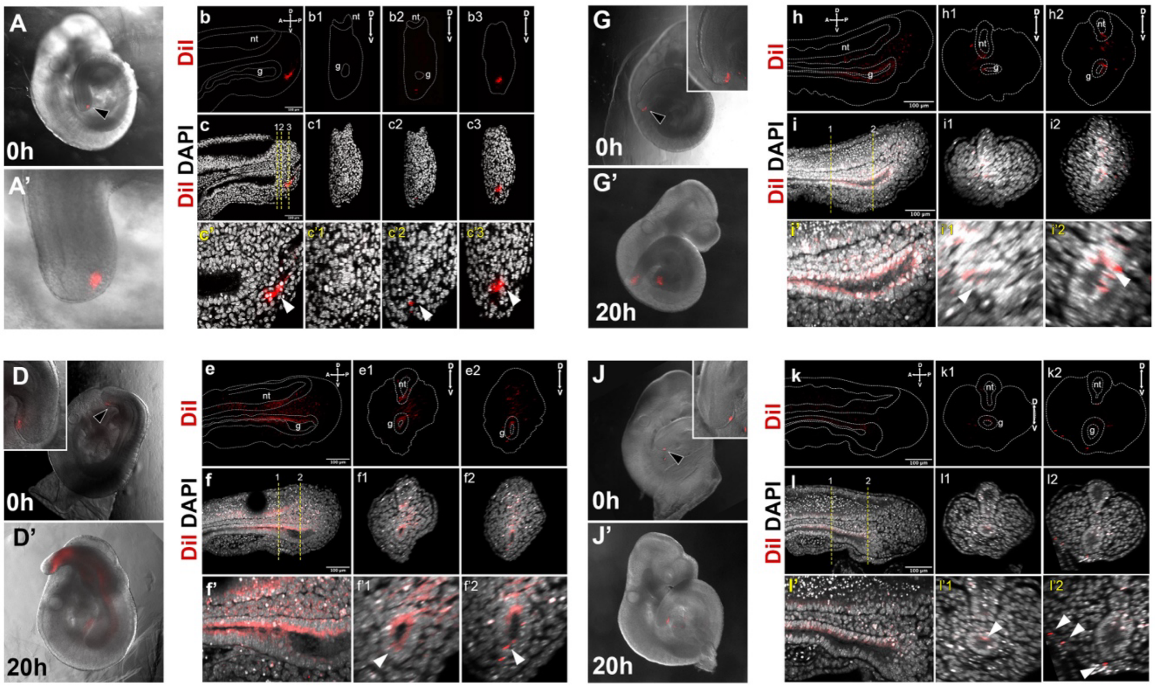


**Supplementary figure 8. DiI labelling of the *Apela*$^{+}$ region in the E9.5 tail bud. A-c3.** Control showing injected region prior to culture. Sagittal and transversal sections shown in b-c3 show absence of DiI label in the gut. **D-i'2.** DiI labelling of E9.5 embryos. Embryos shown in D-i'2 were injected in the region specified in A-c'3. In J-l'2 DiI was injected in tailbud ectoderm (black arrowhead). D, D', G, G', J, J' shown whole mount images of embryos right after the injection (D, G, J) and after 20 h in culture (D', G', J'). Insets in D and G show that gut tube is negative for DiI staining. Inset in J show staining in the ectoderm. e-f'2', h-i'2 and k-l'2 show sagittal (e, f, f', h, i, i', k, l, l') and transversal (e1-f'2, h1-i'2', k1-l'2) optical sections though the tail regions of cultured embryos shown in D', G', J' respectively. e-e2, h-h2, k-k2 DiI labelling, dashed line shows the outline if the tail, neural tube (nt) and tail gut endoderm (g). f-f2, i-i2, l-l2 – overlay of DiI and DAPI channels, lower panels show magnification of stating in the gut. White arrowheads in f'-f'2, i'-i'2 and l'-l'2 show incorporation of DiI-stained cells into gut endoderm. White arrowheads in i'2 show that when ectoderm was injected DiI mainly labels ectoderm, with minor leaking into the dorsal gut (l'1).
